# Genomic insights into six clinical isolates of *Stenotrophomonas maltophilia* from Northern India

**DOI:** 10.1101/2025.01.13.632882

**Authors:** Nazneen Gheewalla, Yuvraj Jadhav, Jaisri Jagannadham, Akansha Tyagi, Sandeep Budhiraja, Bansidhar Tarai, Shraddha Karve

## Abstract

*Stenotrophomonas maltophilia* is a gram-negative opportunistic pathogen that causes respiratory, urinary and bloodstream infections. Due to rising prevalence of difficult to treat *S. maltophilia* infections, it is considered as one of the important pathogens in many regions, including India. Despite its importance in public health, very few studies provide detailed characterization of the genomes of clinical isolates of *S. maltophilia*. In this study, we sequence six isolates of *S. maltophilia* from a tertiary healthcare centre in the Northern India. Along with the culture sensitivity, we report the genomic underpinnings of resistance and virulence in these isolates. In three out of six isolates, we identify a rare beta-lactamase gene *kbl-1* that can confer resistance to variety of beta-lactam antibiotics. Though truncated in two isolates, an intact copy of *kbl-1* is present in the third isolate. This intact copy of *kbl-1* is flanked by insertion elements that are commonly found in several pathogenic species indicating high potential for horizontal gene transfer. To the best of our knowledge, this is a first report of *kbl-1* in *S. maltophilia* from India.

**Importance:** Antimicrobial resistance (AMR) is one of the biggest global public health challenges. Tackling AMR gets complicated as diverse pathogenic species are involved and resistance is manifested against several different kinds of antibiotics. Genomic surveillance of the resistant pathogens is a key to fight this global threat and needs to extend to important emerging pathogens as well. Here we report genomic sequences and associated characteristics of six *Stenotrophomonas maltophilia* isolates from Northern India. Along with several resistance and virulence genes we also discover a rare beta-lactamase, *blaKBL-1*, gene with a high potential of transfer to other species. *blaKBL-1* beta-lactamase confers resistance to several beta-lactam antibiotics like ampicillin, amoxicillin, penicillin G, piperacillin, ceftazidime and cefozopran. This is the first report of the presences of this beta-lactamase from India.

## Introduction

*Stenotrophomonas maltophilia* is a non-fermenting, gram-negative bacterium and one of the important emerging opportunistic, nosocomial pathogens. *S. maltophilia* has been reported to cause respiratory, bloodstream as well as urinary tract infections (Brooke 2012). It causes both community and hospital-acquired infections with majority cases in ICU. (ICMR 2021, 2023). Indian Council for Medical Research (ICMR) classifies drug-resistant *S. maltophilia* as a group III priority pathogen due to the increasing prevalence and difficulty of treating infections (ICMR 2020). *S. maltophilia* is difficult to treat due to its intrinsic resistance to several antibiotics and its ability to form biofilms. (Brooke et al. 2008; Mojica et al. 2022; Sánchez 2015). Together, this results in limited treatment options that mainly include the administration of trimethoprim-sulfamethoxazole or minocycline or levofloxacin or beta-lactam antibiotics such as aztreonam and ceftazidime (as a monotherapy or as a part of combination therapies) (Gibb & Wong 2021; Mojica et al. 2017; Biagi et al. 2020).

*S. maltophilia* is intrinsically resistant to many beta-lactam antibiotics due to the presence of smeABC efflux pump (Li et al. 2002) and beta-lactamases L1 and L2 (Mojica et al. 2019; Lin et al. 2009; Walsh et al. 1994, 1997). Apart from the intrinsic beta-lactamases L1 and L2, *S. maltophilia* may harbor other beta-lactamases such as CTX-M-15, KPC-1, NDM and OXA-48 (Rizi et al. 2024). Kawauchi and colleagues recently reported a novel class A beta-lactamase, KBL-1 in an *S. maltophilia.* This *kbl-1* encoded beta-lactamase had significant activity against several penicillins and weak hydrolytic activity against some cephalosporins. The reported *kbl-1* gene was part of a unique gene cassette that also contained another antibiotic resistance gene (henceforth ‘ARG’) *msrE. kbl-1* and *msrE-*containing regions were flanked by insertion sequence elements IS91 (Kawauchi et al. 2022). IS91 and IS91-like elements have been previously reported in several other pathogenic species (Garcillán-Barcia & De La Cruz 2002) and have a high potential of spreading resistance. Together these studies have shown that genomic surveillance of *S. maltophilia* is important not only for its own sake but also for tracking the resistance elements that can be transferred to other pathogens.

In this study, we report genomic sequences of six *S. maltophilia* isolates from a tertiary healthcare center located in Northern India. We isolated these pathogens during the period of December 2023 to April 2024. We report several resistance and virulence genes that may result in difficult to treat infections. Additionally, we also show that three of the six isolates harbor a rare *kbl-1* beta-lactamase gene. Long-read sequencing of one of the three isolates with *kbl-1* reveals high potential for horizontal gene transfer due to presence of flanking IS elements. To our knowledge, this is the first report from India on the presence of *kbl-1* in *S. maltophilia*.

## Results

### Culture sensitivity and underlying genetic mechanisms

The six *S. maltophilia* isolates that we sequenced belonged to diverse specimen types such as blood, urine and respiratory samples (Table 1). Every isolate was resistant to at least one of the antibiotics tested. Five of the six isolates displayed resistance to trimethoprim-sulfamethoxazole (Table 1, Supplementary table 1). Levofloxacin was the most effective drug with three isolates showing full sensitivity and the remaining isolates showing intermediate sensitivity (Supplementary table 1). Notably, one isolate, SM_6, was completely or partially resistant to all the antibiotics tested (Supplementary table 1).

**Table 1:** Metadata, resistance category and genome assembly information of the six *S. maltophilia* isolates.

| Isolate | Patient age in years, gender, ward | Collection date | Specimen | Resistance observed in culture sensitivity | Sequence type | Genome size (in bp) | Number of coding sequences (cds) | Number of ARGs | Number of virulence genes |
| --- | --- | --- | --- | --- | --- | --- | --- | --- | --- |
| SM_1 | 71, M, ICU | 16/12/2023 | respiratory | Ceftazidime, ticarcillin/ clavulanic acid | ST28 | 4906856 | 4423 | 17 | 81 |
| SM_2 | 5, M, ICU | 19/01/2024 | blood | Chloramphenicol, trimethoprim/ sulfamethoxazole | ST684* | 4566374 | 4088 | 11 | 84 |
| SM_3 | 31, M, ICU | 19/02/2024 | respiratory | Ceftazidime, ticarcillin/ clavulanic acid, chloramphenicol, trimethoprim/ sulfamethoxazole | ST183 | 4848399 | 4380 | 19 | 79 |
| SM_4 | 59, M, ICU | 19/02/2024 | respiratory | Ticarcillin/ clavulanic acid, chloramphenicol, trimethoprim/ sulfamethoxazole | ST183 | 4823937 | 4358 | 19 | 79 |
| SM_5 | 64, M, OPD | 3/04/2024 | urine | Trimethoprim/ sulfamethoxazole | ST621 | 4547968 | 4096 | 11 | 83 |
| SM_6 | 57, M, IPD | 21/04/2024 | urine | Ceftazidime, ticarcillin/ clavulanic acid, trimethoprim/ sulfamethoxazole | ST128 | 4904960 | 4502 | 15 | 93 |

To understand the genomic basis of the observed resistance, we sequenced the entire genomes of these isolates. The genomes had an average size of 4.78 Mb with 4338 coding sequences on an average (Table 1). For each isolate, the number of ARGs ranged from 11 to 19 and the number of virulence genes ranged from 79 to 93. Due to the absence of any reliably detected plasmids (see Methods), we predicted that most of the resistance and virulence genes reside on the chromosome. Apart from genomic basis of observed resistance, we also looked at other ARGs that may confer resistance to antibiotics that were not tested in the culture sensitivity.

We found 26 unique ARGs in the six isolates together. The sme pumps, a group of RND type efflux pumps commonly found in *S. maltophilia* (Li et al. 2002; Zhang et al. 2001), were present in all the isolates. Specifically, smeDEF efflux pump that confers intrinsic resistance in *S. maltophilia* to diverse classes of antibiotics (Zhang et al. 2001; Alonso & Martinez 2000; Brooke 2012), was present in all the six isolates. smeRS, the two-component system which regulates smeABC (Brooke 2012), was present in five of the six isolates. SM_1 also harbored smeABC efflux pump while other isolates, except SM_6, only carried one or more component genes of smeABC (Figure 1 and suppl table 2 for details). Interestingly, the predicted resistance due to the presence of ARGs matched with the culture sensitivity in only some cases. For instance, isolate SM_1 harbored smeABC efflux pump (Figure 1A, supplementary table 2) that can confer resistance to aminoglycosides and a few beta-lactams (Li et al. 2002). Not unexpectedly, SM_1 showed resistance to beta-lactams ceftazidime and ticarcillin/clavulanic acid in the culture sensitivity (Table 1, Supplementary table1). The same isolate also showed multiple gene copies for every component of the smeDEF pump that can confer resistance to tetracyclines, macrolides, chloramphenicol, quinolones and novobiocin (Alonso & Martinez 2000; Zhang et al. 2001; Brooke 2012) (Figure 1A, Supplementary table 2) but remained sensitive to chloramphenicol, levofloxacin and minocycline (Table 1, Supplementary table 1).

**Figure 1.**
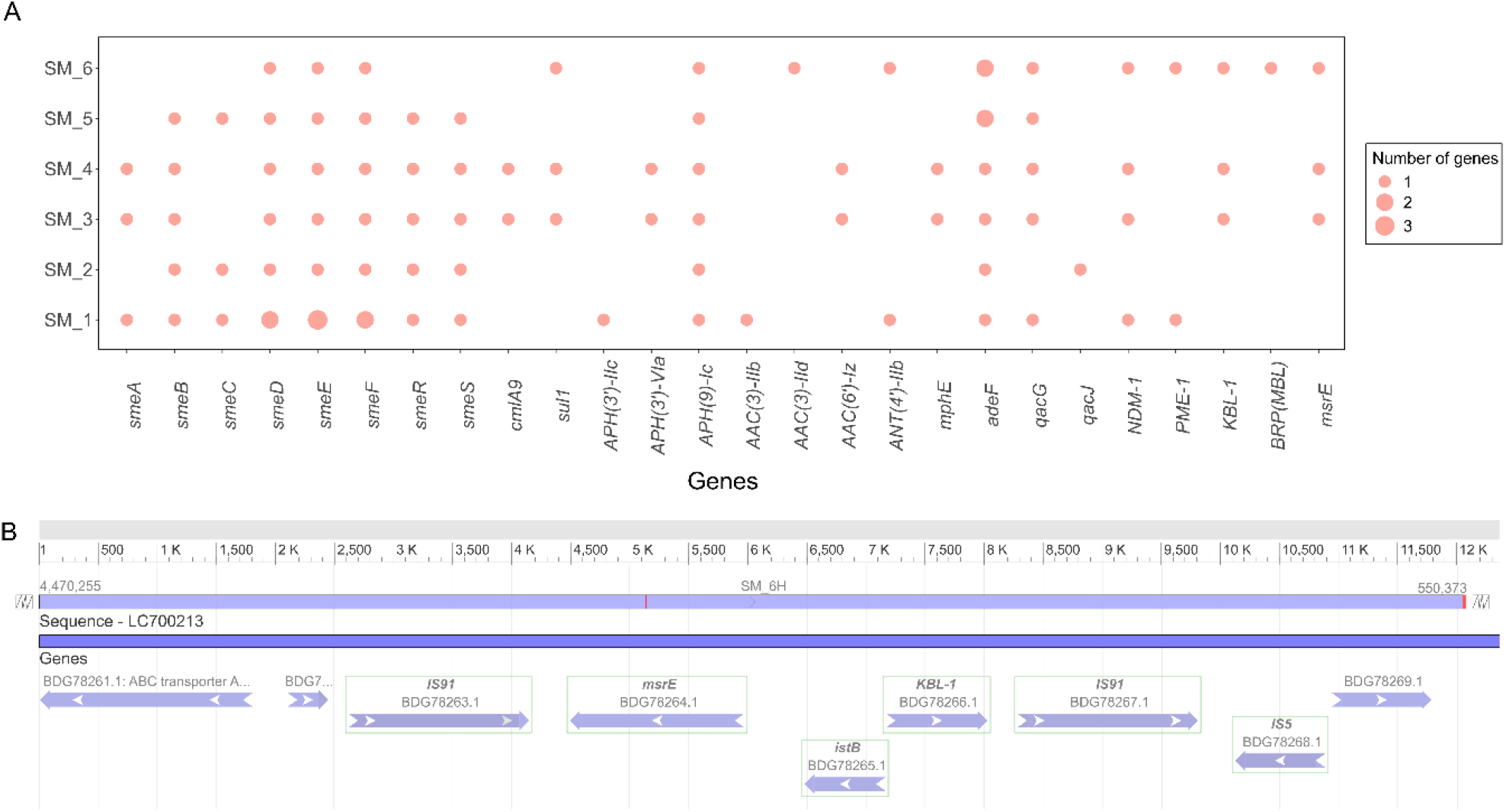
Resistance landscape of *S. maltophilia* isolates. **A:** The panel shows predicted antibiotic resistance genes (ARGs) from the draft genomes of six *S. maltophilia* isolates identified using the CARD-RGI tool. The X-axis indicates different ARGs and the y-axis shows the six isolates. The size of the bubble corresponds to gene copy numbers. The majority of the ARGs are present in a single copy. Three genes that code components of the smeDEF efflux pump, namely *smeD, smeE and smeF*, are present in multiple copies in the isolate SM_1. Similarly, SM_5 and SM_6 harbored two copies of the gene *adeF* that codes for a component of adeFGH efflux pump. **B:** The panel shows a sequence comparison between previously reported gene cassette containing *kbl-1* gene (accession number LC700213.1, (Kawauchi et al. 2022)) and the similar gene cassette from SM_6. The upper horizontal bar in light violet denotes the SM_6 sequence while the lower dark violet bar denotes part of the previously reported cassette. The arrows below denote the components of the gene cassette and their orientation. Part of the previously reported gene cassette depicted here is identical with the same region from SM_6, except a single nucleotide in the *msrE* gene that is indicated by a red line.

*cmlA9*, a gene encoding MFS type efflux pump that confers resistance to chloramphenicol, was also present in two isolates, SM_3 and SM_4 (Figure 1A, Supplementary table 2). Both isolates were resistant to chloramphenicol in the culture sensitivity test as expected (Table 1, Supplementary table 1). Similarly, three isolates, SM_3, SM_4 and SM_6, harbored the *sul1* gene that confers resistance to sulfonamide antibiotics. All three of these show resistance to trimethoprim-sulfamethoxazole, along with SM_2 & SM_5 that did not carry any known sulfonamide resistance conferring gene.

We also found several genes that can confer aminoglycoside resistance in *S. maltophilia. aph(9)-Ic* gene was present in all the six isolates while *aph(3’)-IIc, aac(3)-IIb* and *aac(3)-IId* genes were present in one isolate each. Aminoglycoside resistance-conferring genes *aph(3’)-Via, aac(6)-Iz, ant(4’)-IIb* were present in two isolates each (Figure 1A). For other drug classes, two isolates showed the presence of the *mphE* gene that confers resistance to macrolides. We also detected gene *adeF* which codes for the component of adeFGH efflux pump in all the isolates with two copies in SM_5 and SM_6. adeFGH pump can render *S. maltophilia* resistant to fluoroquinolones and tetracycline (Kaviani et al. 2020; Shahid et al. 2024). Five of the six isolates harbored the gene *qacG* while the remaining isolate carried the *qacJ* gene. *qacJ* and *qacG* genes are known to confer resistance to disinfectants like quaternary ammonium compounds and benzalkonium chloride (Jostein et al. 2003; Heir et al. 1999).

We also looked at the genes conferring resistance to beta-lactams. SM_1, SM_3, SM_4 and SM_6, carried the beta-lactamase gene *ndm-1. SM_1 and SM_6* also harbored another beta-lactamase, *pme-1* and SM_6 additionally carried a metallo-beta-lactamase associated bleomycin resistance gene *BRP* (Figure 1A, Supplementary table 2). Most interestingly, SM_3, SM_4 and SM_6, showed a rare *kbl-1* beta-lactamase gene and closely associated *msrE* gene.

To the best of our knowledge, this is the first report of *kbl-1* gene carrying *S. maltophilia* from India. The *kbl-1* gene is predicted to have evolved from the *psv-1* gene and was first detected in Nepal in an *S. maltophilia* isolate from a case of a blood infection (Kawauchi et al. 2022). The *kbl-1* gene from SM_6 shows complete identity with the reported *kbl-1* gene while it is truncated at the 5’ end in SM_3 and SM_4. Using the completed genome (see methods), we looked at the *kbl-1* and its flanking regions closely. We found that the upstream and downstream regions of the *kbl-1* gene were nearly identical to the previously reported sequence (Figure 1B). An insertion element, *istB*, was present immediately upstream of the *kbl-1* gene, followed by the *msrE* gene. This *msrE-istB-kbl-1* cassette was flanked by two IS91 transposase regions (Figure 1B).

Kawauchi *et al* describe the unique genetic environment surrounding *kbl-1* as a cassette of *orfA-orfB-IS91*-*msrE-istB-kbl-1-IS91-IS5-orfC-orfD-orfE*. In this cassette, orfA is a putative ABC-transporter, orfB is a phosphoglucosamine mutase (Ntn asparginase 2 like protein) and orfs C-E are hypothetical proteins (GenBank: LC700213.1, (Kawauchi et al. 2022)). We find that SM_6 shows a nearly identical cassette of *orfA-orfB-IS91-msrE-istB-kbl-1-IS91-IS5-orfC*. Importantly, the *IS91* flanked ARG-containing regions are identical with a difference of a single SNP in the *msrE* gene (Figure 1B).

#### 2. Virulence genes

*S. maltophilia* is known to carry multiple virulence genes (henceforth ‘VG’s) including genes for secretion, motility, adherence and biofilm formation (Mikhailovich et al. 2024; Bhaumik et al. 2023; Trifonova & Strateva 2019). We detected several VGs in every isolate. We found 138 unique VGs in all six isolates together with number of unique VGs in a single isolate ranging from 79 to 93 (Supplementary table 3). Most prevalent VGs were associated with motility or chemotaxis such as genes from the *flg, fla* and *fli* gene families (Liu et al. 2022). Apart from these other genes related to motility or chemotaxis were also present. Examples are *motA, motB, motD, cheA, cheR* and *cheW* (Liu et al. 2022). All isolates harbored genes encoding proteins of effector delivery systems such as the *gsp* genes associated with the bacterial type II secretory system. The biofilm-associated genes, *algA, algC* and *mucD* were also present in all the isolates, while SM_6 additionally harbored the *algW* gene (Liu et al. 2022). Genes associated with pili, such as *pilB, pilG, pilJ* and *pilR*, were also present in every isolate (Liu et al. 2022).

## Discussion

We sequenced six *S. maltophilia* isolates from a tertiary health care center and characterized their resistomes and virulomes. We detected several resistance and virulence genes on the chromosome. Interestingly, we did not detect any plasmid in any of the isolates. One reason for this absence could be the lack of well-annotated plasmids in the database from *S. maltophilia*. However, completed genome of SM_6 showed a single genomic contig supporting the absence of any plasmids.

We also found a few cases of mismatch between the predicted (by CARD-RGI tool) and observed (by culture sensitivity testing) resistance patterns. While some of them could be genuine cases of disagreement, others may be due to truncated genes or lack of gene expression or some other unidentified factor. Detailed look at every instance of disagreement was out of scope of this work but can be one of the promising future direction.

Three of the six isolates harbored the rare *kbl-1* gene that has previously only been reported in one *S. maltophilia* isolate in Nepal. In one of the isolates, intact *kbl-1* gene along with the *msrE*, is flanked by transposable insertion elements IS91 forming a unique gene cassette. IS91 elements are found in pathogenic *E. coli* and IS91-like elements are found in many other pathogenic species such as *Pseudomonas, Klebsiella, Shigella, Salmonella, Vibrio* (Garcillan-Barcia M et al 2002). This suggests the potential for horizontal transfer across pathogenic species. Our study is the first report of *kbl-1* containing *S. maltophilia* from India and underlines the importance of genomic surveillance for *S. maltophilia*.

## Materials and methods

We selected six *S. maltophilia* isolates with distinct antibiotic resistance profiles in the culture sensitivity test. We stored the selected isolates as glycerol stocks for genomic DNA extraction. For the glycerol stock preparation, we inoculated overnight grown cultures in tryptone soy broth with 15% glycerol (HiMedia LQ278II) and stored at −80°C. Before DNA extractions, we revived part of this glycerol stock on McConkey agar (Himedia MH081) plates and resuspended the revived culture in water for injection. We used the Qiagen blood and tissue kit (Cat no: 69504) as per the accompanying protocol to extract the genomic DNA from this cell suspension. We eluted the purified genomic DNA with 10mM of Tris-Cl, pH 8 (HiMedia, ML013) and measured the concentration of the eluted genomic DNA using Qubit (Thermo, Q33231). We then sent the extracted genomic DNA for short-read sequencing using the NovaSeq 6000 Illumina platform at the National Center for Biological Sciences, Bangalore, India. We also sent one isolate, SM_6, for long-read sequencing using the Oxford Nanopore Technology (ONT) at the CSIR-Centre for Cellular and Molecular Biology at Hyderabad, India where they used the PRO114M (R10.4.1) flowcell on a PromethION 24 for sequencing the one *S. maltophilia* isolate.

We built *de novo* short-read assemblies for all six isolates using a customized pipeline (Karthikeyan). We then used the CARD-RGI tool v6.0.3 (database version 3.3.0) (accessed on 3^rd^ December 2024) (Alcock et al. 2023) to detect ARGs in the assembled genomes. We retained ‘perfect’ and ‘strict’ matches for identifying ARGs but nudged the loose hits with a match greater than 95% to the ‘strict’ category. We checked the presence of virulence genes using the BLASTn v2.9.0 (Altschul et al. 1990) and the full set of DNA sequences from the VFDB database (Liu et al. 2022). We ignored any duplicate entries, if present, and marked the virulence gene as present when at least one match was found. We also performed MLST (Multi Locus Sequence Typing) analysis (v2.0.9) (Bartual et al. 2005; Camacho et al. 2009; Griffiths et al. 2010; Jaureguy et al. 2008; Larsen et al. 2012; Ludovic et al. 2004; Wirth et al. 2006).

For SM_6 isolate, we also built the hybrid assembly with a long-read first approach using a customized pipeline (Karthikeyan). We used BLASTn webtool (accessed on 27th December 2024) (Altschul et al. 1990) with default thresholds to compare the sequence of *kbl-1* in assemblies from the current study (both short-read and hybrid) and the reported *kbl-1* gene (accession LC579778.1). We used hybrid assembly to study the similarity of the *kbl-1* gene and its flanking regions with the previously reported *kbl-1* flanking region submitted by Kawauchi et al (accession LC700213.1) (Kawauchi et al. 2022).

For the six short-read assemblies and one hybrid assembly, we used MobSuite (v3.1.9) (Robertsen and Nash, 2018, Robertsen et al 2020) to detect plasmids. The software reported a few putative plasmids but did not provide the replicon and relaxase type for those. As a result, we could not confirm the presence of any plasmids.

## Data Availability

Complete genome of the SM_6 hybrid assembly and illumina sequencing based sequence reads (sra) of the six isolates are submitted to NCBI under Bioproject accession number PRJNA1209692.

## Ethics statement

Institutional ethics committee approval for the multi-center study was obtained at Ashoka University. Additionally, the clinical partner, Max Healthcare, obtained ethical clearances from their ethics committee.

## Acknowledgement

We acknowledge Rockefeller Foundation Grant Number 2021 HTH 018 and financial aid by Axis Bank for funding the multi-centre AMR mapping and genomic analysis study at Ashoka University. We thank technical team from Max Healthcare for maintenance of pathogen culture stocks and DNA extractions. We thank Vasundhara K. for discussions on the bioinformatic analysis framework. We thank Dr. Aradhita Baral for the project management support.

## Author credits

NG and SK conceived the study. AT, BT and SB obtained isolates and associated metadata, culture sensitivity report and genomic DNA. NG analysed the metadata. YJ built the de novo assemblies from the raw sequence data. NG, YJ and JJ performed bioinformatic analysis. NG & SK prepared the manuscript. SK edited the final manuscript.

